# Increased amphetamine-induced hyperactivity in *translin (Tsn)* KO mice is driven by elevated adiposity

**DOI:** 10.1101/2020.10.08.331447

**Authors:** Xiuping Fu, Aparna P. Shah, Jacqueline Keighron, Ta-Chung M. Mou, Bruce Ladenheim, Jesse Alt, Daisuke Fukudome, Minae Niwa, Kellie Tamashiro, Gianluigi Tanda, Akira Sawa, Jean-Lud Cadet, Rana Rais, Jay M. Baraban

**Affiliations:** Solomon H. Snyder Department of Neuroscience, Department of Psychiatry and Behavioral Sciences, John Hopkins Drug Discovery, Department of Neurology, Department of Biomedical Engineering, Johns Hopkins School of Medicine, Baltimore, MD 21205; Department of Mental Health, Johns Hopkins University Bloomberg School of Public Health, Baltimore MD 21205; Department of Psychiatry and Behavioral Neurobiology, University of Alabama at Birmingham, Birmingham, AL 35233; Medication Development Program, National Institute on Drug Abuse, Intramural Research Program, National Institutes of Health, 333 Cassell Drive, Baltimore, MD, 21224; Molecular Neuropsychiatry Research Branch, Intramural Research Program, NIDA/NIH/DHHS, 251 Bayview Boulevard, Baltimore, MD, 21224

## Abstract

The translin/trax microRNA-degrading enzyme mediates activity-induced changes in translation that underlie several long-lasting forms of cellular plasticity. As translin and trax are expressed in dopaminergic and striatal neurons, we proceeded to investigate whether deletion of *Tsn* blocks amphetamine sensitization, a long-lasting, translation-dependent form of behavioral plasticity,

Although we expected constitutive *Tsn* deletion to impair amphetamine sensitization, we found, instead, that it enhances the hyperlocomotion produced by the initial dose of amphetamine. Since these mice display elevated adiposity, which alters pharmacokinetics of many drugs, we measured brain amphetamine levels in *Tsn* knockout mice and found that these are elevated. We also found that diet-induced increases in adiposity in WT mice correlate with elevated brain amphetamine levels. As amphetamine and its analogues are widely used to treat attention deficit disorder, which is associated with obesity, further studies are needed to assess the impact of adiposity on amphetamine levels in these patients.

## Introduction

Recent studies have focused attention on the role of the microRNA system in the pathophysiology of psychiatric diseases [1–4]. However, our understanding of how microRNA signaling pathways operate normally in the nervous system is quite limited, making it difficult to assess how altered microRNA signaling may contribute to these disorders. To help elucidate the physiological function of microRNA signaling in brain, we are investigating the role of the translin (TN)/trax (TX) microRNA-degrading enzyme, an RNase that targets a small population of microRNAs [5, 6]. We are focusing on this enzyme because expression of TN and TX is enriched in brain, where it is localized preferentially to neurons [7].

Characterization of *translin (Tsn)* knockout (KO) mice, which lack the TN/TX complex, has enhanced our understanding of the role of this microRNA-degrading enzyme *in vivo*. These studies have revealed several prominent phenotypes displayed by *Tsn* KO mice: blockade of several forms of long-lasting synaptic plasticity [8, 9], decreased anxiety [10], protection from vascular stiffening [11], and development of robust adiposity in the context of normal body weight [12]. Furthermore, defining the mechanism mediating these phenotypes suggests that TN/TX operates in the following manner: TN/TX activation by cellular stimulation triggers rapid degradation of a small population of microRNAs. Degradation of these microRNAs reverses silencing of their target mRNAs, allowing *de novo* protein translation to proceed. Thus, the TN/TX complex appears to be well positioned to mediate translation-dependent forms of cellular plasticity.

In pilot studies, we noted that TN and TX are expressed in midbrain dopaminergic (DA) neurons and striatal neurons, among other brain regions. These neuronal populations play key roles in mediating the action of psychostimulants [13–15], including sensitization, a long-term form of behavioral plasticity that is dependent on translation and thought to underlie the reinforcing properties of these drugs [16–18] Furthermore, recent studies have yielded compelling evidence demonstrating that the microRNA system plays a prominent role in mediating behavioral responses to these drugs [19–23]. Accordingly, we hypothesized that the TN/TX RNase mediates this form of plasticity and proceeded to check the locomotor response to amphetamine in *Tsn* KO mice.

## Methods and Materials

### Animals

All mice were housed in ventilated racks and maintained on a 14-h/10-h light/dark cycle with access to regular chow (2018SX Teklad Global, Frederick, MD) and water *ad libitum*. All procedures were performed in accordance with the NIH Guide for the Care and Use of Laboratory Animals and approved by the Johns Hopkins Animal Care and Use Committee.

The *Tsn* KO mice used in these studies are from the line generated in Dr. Kasai’s laboratory [24] and provided by the JCRB Laboratory Animal Resource Bank of the National Institute of Biomedical Innovation (translin KO: Nbio055).

Mice with floxed alleles of *Tsn* or *Tsnax* were generated on a C57BL/6J background using CRISPR/Cas9 technology as described in Fu et al.[25]. The following Cre driver or reporter lines were obtained from JAX labs or MMRRC (Drd1-Cre (EY217); Ai9 #007909; DAT-Cre #006660; Ubiquitin C-CreERT2 #007001). Genotyping was performed on tail snips by Transnetyx, Inc. (Cordova, TN).

To examine the effects of elevated adiposity in WT mice (C57BL6/J), we separated 3 month old mice obtained from JAX labs into three groups as described by Fordahl et al. [26]: regular chow, unrestricted access to HF diet (D12492; Research Diets, New Brunswick, NJ), and limited access to the same HF diet. The limited access mice had their feed changed from regular chow pellets to the HF pellets for a 2-hour period on Monday, Wednesday and Friday. Mice were kept on this regimen for 7 weeks. All these mice underwent scanning to determine their adiposity (% body fat) 5 days after being switched back to regular chow. After being scanned for body composition, mice underwent either open field locomotor testing or collection of plasma and brain samples that were used for determination of amphetamine levels.

### Immunostaining

Three to four-month old male mice were deeply anesthetized with chloral hydrate (400 mg/kg, i.p.) for transcardial perfusion with 4% paraformaldehyde in 0.1 M phosphate-buffered saline pH 7.4 (PBS). Their brains were dissected and stored in 4% paraformaldehyde for at least 24 h, and then washed with PBS and transferred to 30% sucrose in PBS. After the brains totally sank, 30 μm sections were cut with a freezing microtome.

For TN immunostaining, sections underwent heat-induced antigen retrieval by incubating them in 10mM NaCitrate for 30 min at a temperature of 70°C. After the sections cooled down to room temperature, they were washed with PBS three times, blocked with 3% BSA and 0.1% Triton X-100 in PBS for 1h, and then incubated with primary antibodies against TN [27] and tyrosine hydroxylase (TH; MilliporeSigma, Burlington, MA) overnight at 4°C. After three washes, the sections were incubated with Alexa Fluor 488 goat anti-rabbit (Jackson ImmunoResearch Laboratories, West Grove, PA) and Alexa Fluor 594 goat anti-mouse (Jackson ImmunoResearch Laboratories) in 1.5% Normal Goat Serum (Vector Laboratories, Burlingame, CA) in PBS for 1h at room temperature. After three additional washes with PBS, the sections were mounted on slides, and coverslipped.

For TX immunostaining, sections were washed with PBS three times and blocked with 3% BSA and 0.1% Triton X-100 in PBS for 1h, followed by incubation with primary antibodies against TX [28] overnight at 4°C. After three washes, the sections were incubated with biotinylated goat anti-rabbit (Vector Laboratories) in PBS for 1h at room temperature. After three washes with PBS, the sections were incubated with AB solution (ABC kit, Vector Laboratories) for 90 min. Sections were washed with TBS three times, and incubated with tyramide (TSA kit, PerkinElmer) for 10 min. After one wash with TBS and three washes with PBS, sections were mounted on slides, and coverslipped.

All images were obtained with a Zeiss LSM 800 confocal microscope.

### Western Blotting

Forebrain, nucleus accumbens and dorsal striatum tissue samples were homogenized in RIPA buffer (Cell Signaling Technology, Danvers, MA) containing a cocktail of protease and phosphatase inhibitors (MilliporeSigma). The concentration of total protein was determined using the Pierce BCA Protein Assay Kit (Thermo Fisher Scientific, Waltham, MA). Equal amounts of total protein were separated electrophoretically on an SDS-PAGE gel, transferred to a PVDF membrane (Bio-Rad, Hercules, CA) and immunoblotted with antibodies: anti-DAT (MilliporeSigma), anti-D2R (MilliporeSigma), anti-TH (MilliporeSigma), and anti-tubulin (Cell Signaling Technology), anti-TN and anti-TX. Blots were developed with the ECL system (Thermo Fisher Scientific). Band intensities were quantified from digital images by densitometry using ImageJ.

### Open-Field Locomotor Activity

Changes in locomotor activity in response to amphetamine were assessed in an open field arena. 3 to 4-month old male mice were first allowed to habituate to the arena (50 × 50 cm) for 30 minutes. Following habituation to the context, mice were given an i.p. injection of saline and then placed back in the center of the open field and allowed to explore the arena for another 30 minutes. At this point, mice were given an intraperitoneal (i.p.) injection of D-amphetamine hemisulfate (2.5mg/kg of total body weight or 3.5 mg/kg of lean mass; MilliporeSigma), placed back in the arena and then monitored for an additional 40-50min.

### Tissue Dopamine Levels

Mice were sacrificed, their brains removed and placed on ice. Dorsal striatum and nucleus accumbens were dissected and weighed, then flash frozen and stored at - 80°C. Tissue samples were ultrasonicated in 0.1 M perchloric acid, and stored at −80°C until extraction. Upon thawing, the samples were homogenized in 0.1 M perchloric acid and centrifuged at 25,000 g for 12 minutes. Dopamine levels were measured by HPLC with electrochemical detection. The analytical column was a SunFire C18 5 lm (4.6 · 150.0 mm) from Waters (Milford, MA, USA). The mobile phase was 0.01 M sodium dihydrogen phosphate, 0.01 M citric acid, 1.2 mM sodium EDTA, 1.2 mM sodium 1-heptane sulfonic acid, 10% methanol, pH 3.5; the flow rate was 1.0 mL/min and the column temperature was 34°C. The installation consisted of a Waters 717 Plus automated injection system, a Waters 1525 Binary pump, and an ESA Coulochem III detector (Dionex, Sunnyvale, CA, USA). Waters Breeze system was used for analysis.

### Body Composition

Body composition was determined by using a nuclear magnetic resonance scanner (EchoMRI-100, Houston, TX).

### Plasma and Brain Amphetamine Levels

Amphetamine levels in plasma and brain samples were measured using high-performance liquid chromatography with tandem mass spectrometry (LC/MS-MS). Methanol, containing 0.5μM losartan as an internal standard, was used to extract amphetamine from plasma and brain. Standards were prepared by spiking amphetamine in naïve mouse tissue from 0.01 - 100 μmol/g in a half log dilution series. Plasma (20μL) and brain samples and corresponding standards were placed in low retention microfuge tubes with 5 μL/mg of extraction solution for protein precipitation. Samples were vortex mixed, followed by centrifugation at 16,000 x g for 5 minutes at 4°C. The supernatants (80 μL) were transferred to a 96 well plate and 2 μL was injected for analysis. Samples were analyzed on an UltiMate 3000 UHPLC coupled to Q Exactive Focus Orbitrap mass spectrometer (Thermo Fisher Scientific). Samples were separated on an Agilent Eclipse Plus C18 RRHD (1.8 μm) 2.1 × 100 mm column. The mobile phase consisted of water + 0.1% formic acid (A), and acetonitrile + 0.1% formic acid (B). Separation was achieved at a flow rate of 0.4 mL/min using a gradient run, from 97.5/2.5 (A/B) to 5/95 (A/B) over 1.5 minutes, maintaining at 5/95 (A/B) for 1 minute, and then re-equilibrating for 1 minute. Samples were introduced to the source through heated ion spray with the capillary temperature setting at 350° C and spray voltage of 3.5 kV. Nitrogen was used as the sheath and auxiliary gas with the settings of 30 and 5 respectively. Quantification was performed in product-reaction monitoring (PRM) mode with collision energy setting of 10. Transitions 136.1121 m/z to 119.0876 m/z and 91.0563 m/z (amphetamine) and 423.1695 to 377.1522 m/z and 207.0915 m/z (losartan) were monitored. Data were acquired and quantified with Xcalibur software.

### Tamoxifen Treatment

Tamoxifen (10mg/ml in corn oil; MilliporeSigma) was administrated daily (100 mg/kg, i.p.) for 6 consecutive days to Tsn ^fl/fl^ and UBC-Cre ^+/-^/Tsn ^fl/fl^ mice. Three weeks later, these mice were used to monitor the locomotor response to amphetamine administration.

### Statistical Analysis

Data are presented as mean ± SEM. Statistical significance was evaluated using GraphPad Prism 7 (GraphPad Software, La Jolla, CA). Student’s t-test was used to compare groups in single variable experiments. Repeated measures (RM) two-way ANOVA was used to analyze multiple variable experiments. Pairwise comparisons were made using Bonferroni’s post-hoc test. Differences were considered significant at p<0.05.

## Results

### TN and TX are expressed in midbrain DA and striatal neurons

In prior immunostaining studies characterizing the expression of TN and TX in hippocampus and cortex [7], we noticed prominent neuronal staining in striatum and several brain stem areas. To check if TN and TX are expressed in midbrain DA neurons, we performed double immunostaining for TH and either TN or TX. These studies demonstrate clear co-expression of TN and TX in DA neurons (Figure 1A). TX staining in striatum is present in both D1R-positive and D2R-positive neurons, as we found a high degree of co-expression of TX in neurons stained for DARPP-32, which is expressed in both populations [19]. Furthermore, we conducted TX staining in tdTomato reporter mice (Ai9) that express this marker selectively in D1R-positive neurons, under the control of Drd1-Cre. We found that TX is expressed in both D1R-positive neurons, which are labeled with tdTomato, as well as in presumptive D2R-positive neurons, which are not (Figure 1B).

**Figure 1:**
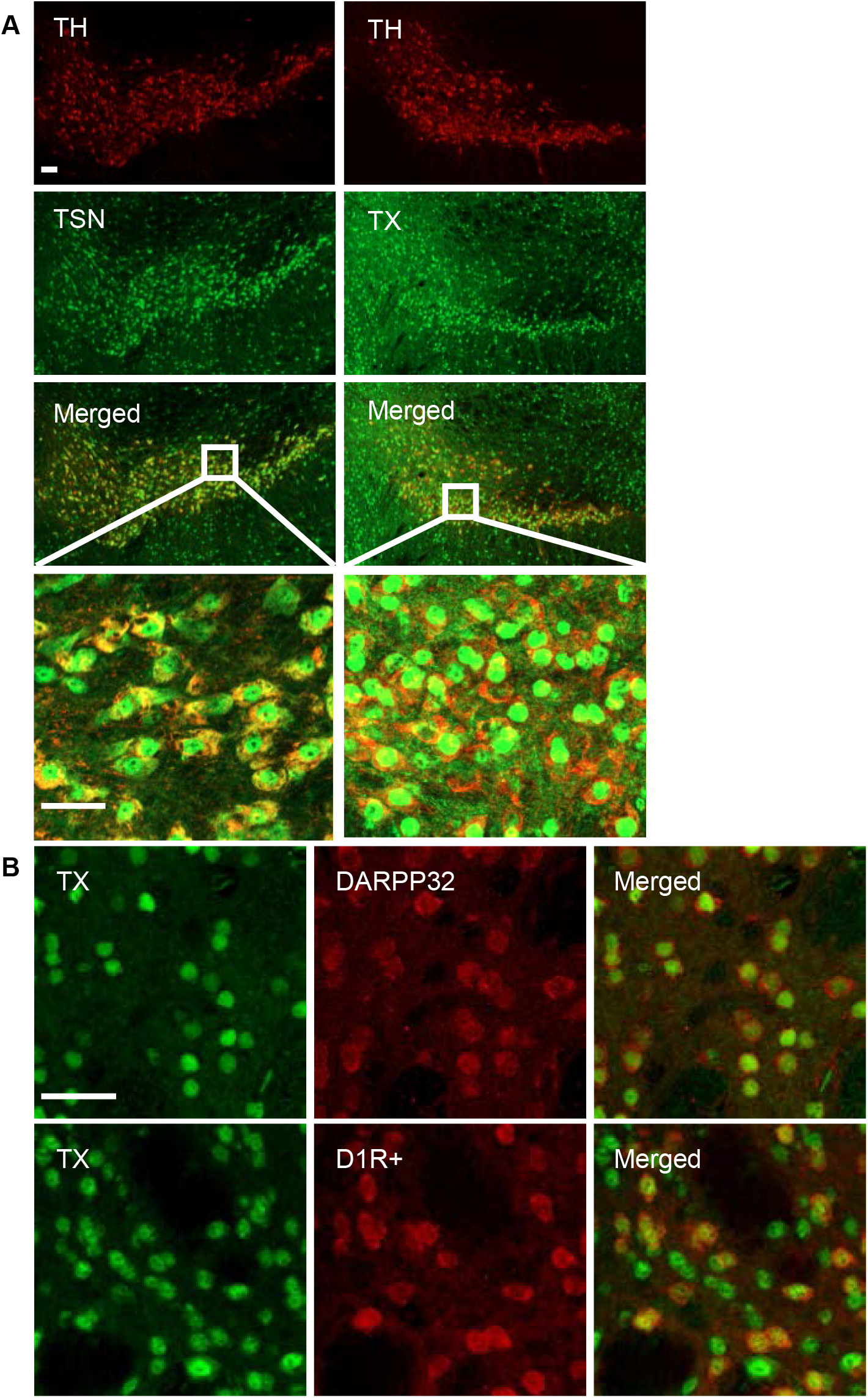
Localization of TN and TX to DA neurons and striatal neurons. (A) Double immunostaining of midbrain sections for TH (red) and TN or TX (green) shows TN or TX expression located in the VTA or substantia nigra. (B) Immunostaining in striatal medium spiny neurons shows prominent nuclear TX staining (green) in DARPP-32 positive neurons (red) (top row). TX staining of sections from mice expressing tdTomato in D1R+ neurons demonstrates that TX is expressed in both D1R-positive neurons as well as in D1R-negative neurons (bottom row). Scale bar, 30 μm.

### Tsn KO mice display increased locomotor response to amphetamine

To investigate the role of the microRNA system in regulating behavioral responses to amphetamine, we compared its ability to stimulate open field locomotor activity in *Tsn* KO mice and WT littermates (Figure 2A). Although we had hypothesized that *Tsn* deletion might impact responses elicited by repeated amphetamine administration, such as locomotor sensitization, which are mediated by changes in protein translation [17], we found, unexpectedly, that the locomotor response to the first injection of amphetamine is markedly enhanced in *Tsn* KO mice. In light of this observation, we checked if *Tsn* deletion impacts the midbrain DA pathway, which plays a major role in mediating the behavioral effects of amphetamine [14, 15].

**Figure 2:**
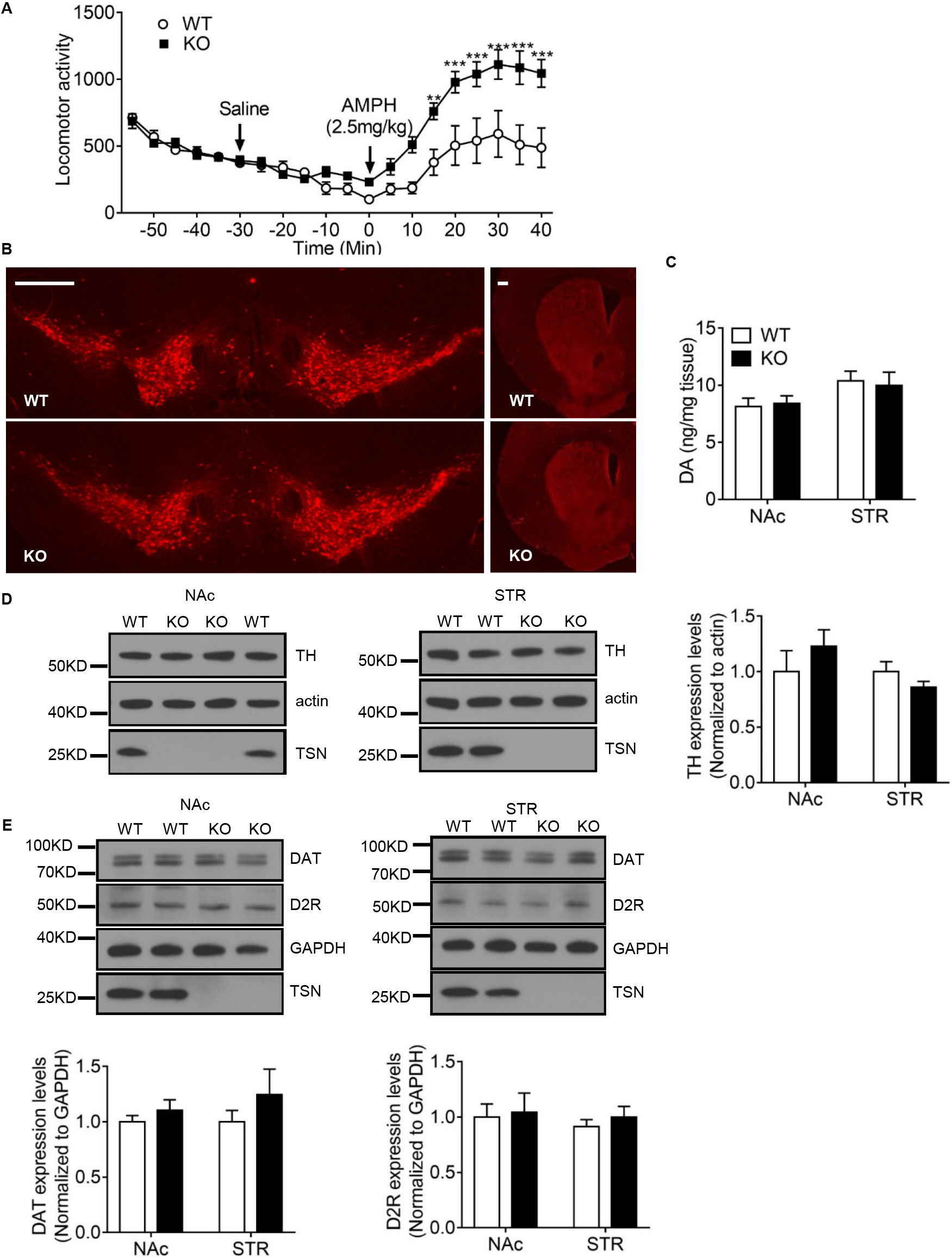
*Tsn* KO mice display increased locomotor response to amphetamine (AMPH) but normal levels of tissue DA, TH, DAT and D2R. (A) Amphetamine-induced (2.5 mg/kg, i.p.) locomotor activity is increased in KO mice. Locomotor activity was monitored every 5 minutes and arrowheads indicate time of injections. n = 8/group. Two-way ANOVA with RM revealed a significant effect of time (p<0.0001), a significant effect of genotype (p=0.0095), as well as a significant interaction between time X genotype (p<0.0001). Bonferroni post-hoc testing was done to identify individual time points that were significantly different. **p<0.01, and ***p<0.001. (B) Immunostaining shows comparable TH staining of DA neurons (left panel) and striatal terminals (right panel) in WT and translin KO mice. Scale bar, 500 μm. (C) Tissue DA levels in the nucleus accumbens (NAc) and striatum (STR) are similar in WT and translin KO mice. n = 5-6 /group. (D and E) Immunoblotting shows normal TH, DAT, D2R expression levels in NAc and STR in translin KO mice. Image J was used for quantification. n = 6/group. Data are expressed as mean ± SEM. Statistical significance was assessed by Student’s t-test.

Qualitatively, staining of midbrain and striatal sections from WT and *Tsn* KO mice for tyrosine hydroxylase (TH), showed a comparable pattern of expression of this enzyme in dopamine neuronal cell bodies and striatal projections between the two genotypes (Figure 2B). To obtain a quantitative assessment of the status of dopamine neurons, we measured tissue dopamine levels in nucleus accumbens (NAc) and dorsal striatum (STR) by HPLC (Figure 2C), as well as protein levels of TH (Figure 2D), dopamine transporter (DAT) and D2 dopamine receptor (D2R) by immunoblotting (Figure 2E), in both the NAc and STR. We did not detect any differences between samples obtained from *Tsn* KO mice and WT littermates in these assays.

### Increased brain levels of amphetamine in Tsn KO mice: correlation with increased adiposity

As *Tsn* KO mice display robust adiposity and adiposity impacts the pharmacokinetics of many drugs [29–31], we considered the possibility that their enhanced locomotor response to amphetamine might be due to increased brain amphetamine levels. Since the amphetamine dose (2.5 mg/kg) used in monitoring locomotor response is in the mid-portion of the ascending part of the dose-response curve [32], increased levels of amphetamine in *Tsn* KO mice could account for their enhanced response in this assay. Consistent with this scenario, we noted a positive correlation between locomotor response to amphetamine and adiposity (%fat mass) that is pronounced in WT mice (Figure 3A), but also present when including both WT and *Tsn* KO mice. To test this directly, we collected both plasma and brain samples at 30 minutes following administration of amphetamine (2.5 mg/kg, i.p.). We found that brain levels of amphetamine are increased in *Tsn* KO mice, with a strong correlation between adiposity and amphetamine levels in either brain or plasma (Figure 3B and 3C). Thus, these findings: 1) indicate that the increased locomotor response to amphetamine in *Tsn* KO mice reflects increased brain levels of this drug, and 2) suggest that the alteration in amphetamine pharmacokinetics is related to the elevated adiposity, or other correlated metabolic abnormalities, displayed by *Tsn* KO mice.

**Figure 3:**
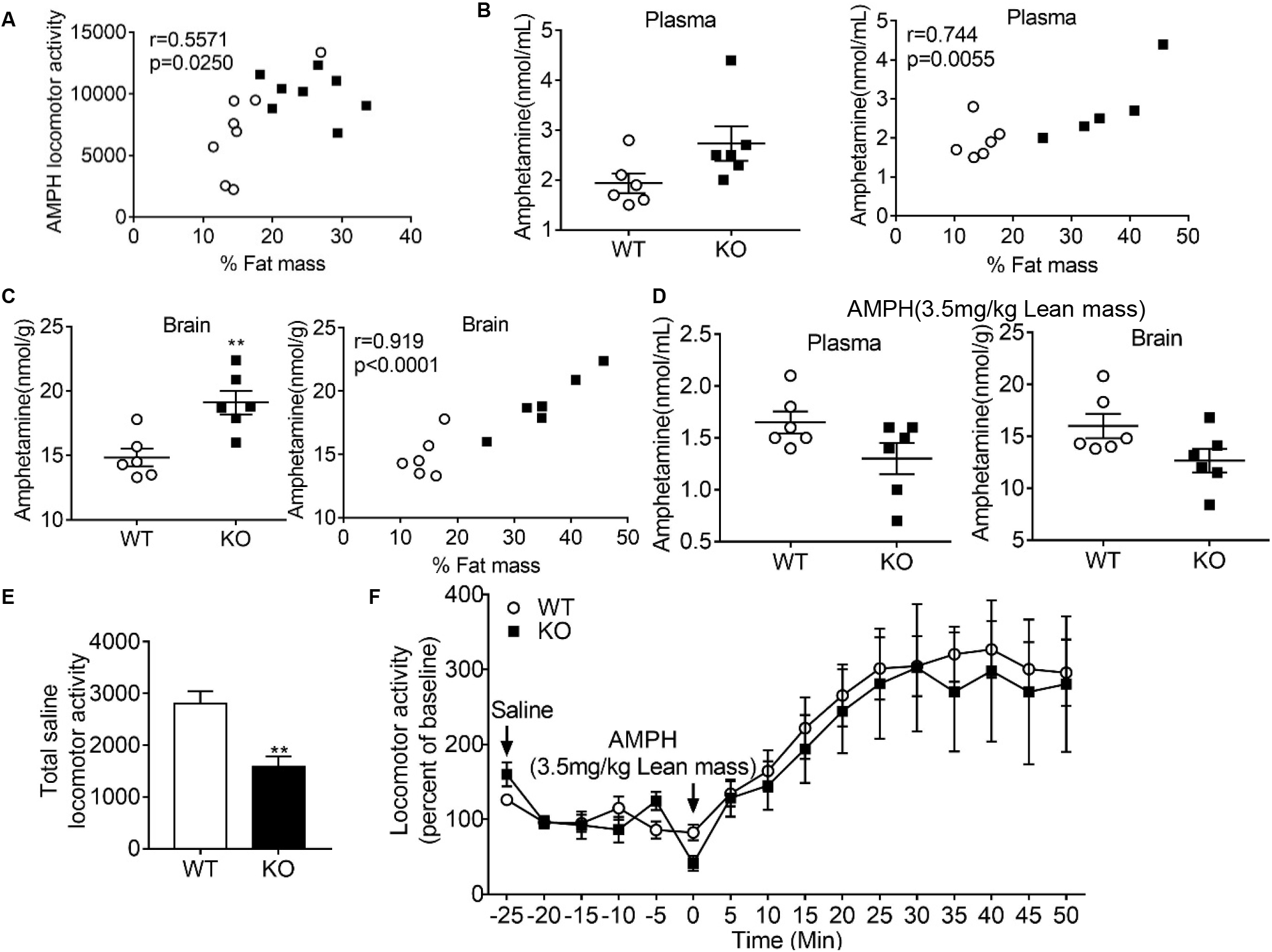
Correlation of fat mass with AMPH-induced locomotor activity and AMPH levels in plasma and brain. (A) The total locomotor activity induced by AMPH (2.5 mg/kg i.p.) was correlated with % fat mass (r = 0.5571, P =0.025). n= 8/group. (B) Plasma AMPH levels (in nmol/mL) (left), and correlation with % fat mass (right; r =0.744, P =0.0055). 2.5mg/kg AMPH was injected i.p. based on body weight. n = 6/group. (C) Brain AMPH levels (in nmol/g) (left), and correlation with % fat mass (right; r =0.919, p<0.0001). 2.5mg/kg AMPH was injected i.p. based on body weight. n = 6/group. (D) Plasma and brain AMPH levels when 3.5 mg/kg AMPH was injected i.p. based on lean mass. n=6/group. (E) Prior to i.p. injection of 3.5 mg/kg based on lean mass, baseline locomotor activity monitored following saline injection was lower in Tsn KO mice than in WT littermates. (F) The normalized locomotor response to AMPH shown as a percentage of the baseline level (100%) is comparable between groups. n=7-8/group. Data are expressed as Mean ± SEM. Statistical significance indicated by asterisks was assessed by Student’s t-test. **p<0.01.

Since *Tsn* KO mice exhibit a prominent increase in adiposity (~2-3 fold) without an increase in total body weight [12], both WT and *Tsn* KO mice received comparable amounts (2.5 mg/kg of total body weight) of amphetamine in experiments monitoring its effects on locomotor activity. On the other hand, *Tsn* KO mice received a higher dose with respect to their reduced lean body weight. As lipophilic compounds, such as amphetamine, are preferentially distributed to tissues with high blood flow, such as brain [33], it is conceivable that amphetamine would be preferentially delivered to brain at the expense of adipose tissue, which has a very low perfusion rate. These findings suggest that calculating the amount of amphetamine administered to each mouse based on their lean body mass, rather than total body weight, should eliminate the disparity in brain amphetamine levels between WT and *Tsn* KO mice. To test this inference, we administered a dose of 3.5 mg/kg amphetamine, ip, based on lean body weight, and collected plasma and brain samples 30 minutes later. Consistent with this hypothesis, this modified dosing regimen produced similar levels of amphetamine in plasma or brain of WT and *Tsn* KO mice (Figure 3D), and elicited comparable locomotor responses to amphetamine in both groups. Since this cohort of *Tsn* KO mice displayed lower baseline activity prior to administration of amphetamine (Figure 3E), we plotted the post-amphetamine locomotor activity as a percentage of baseline activity (Figure 3F).

### Increased adiposity in WT mice elevates brain amphetamine levels

Although these findings indicate that the robust adiposity present in *Tsn* KO mice elicits elevated brain amphetamine levels, they do not rule out the possibility that the elevated brain amphetamine levels detected in *Tsn* KO mice could be due to other effects of *Tsn* deletion that are correlated with their elevated adiposity. Accordingly, to exclude this potential confound, we assessed the effects of diet-induced adiposity on amphetamine brain levels in WT mice. WT mice were separated into three groups: one group was maintained on regular chow (Chow), one group was switched to a HF diet (HF), and the third group had limited access to the HF diet (LMHF; 2 hours on MWF). After seven weeks on these regimens, mice were scanned to determine their %body fat, and then used to assay locomotor response to amphetamine or measure amphetamine levels in plasma and brain. Comparison of the baseline activity levels following administration of saline revealed that mice in the HF group had reduced activity compared to the LMHF mice (Figure 4A). Analysis of locomotor activity following amphetamine administration demonstrated that the HF mice exhibited an elevated response to amphetamine compared to the other two groups (Figure 4B).

**Figure 4:**
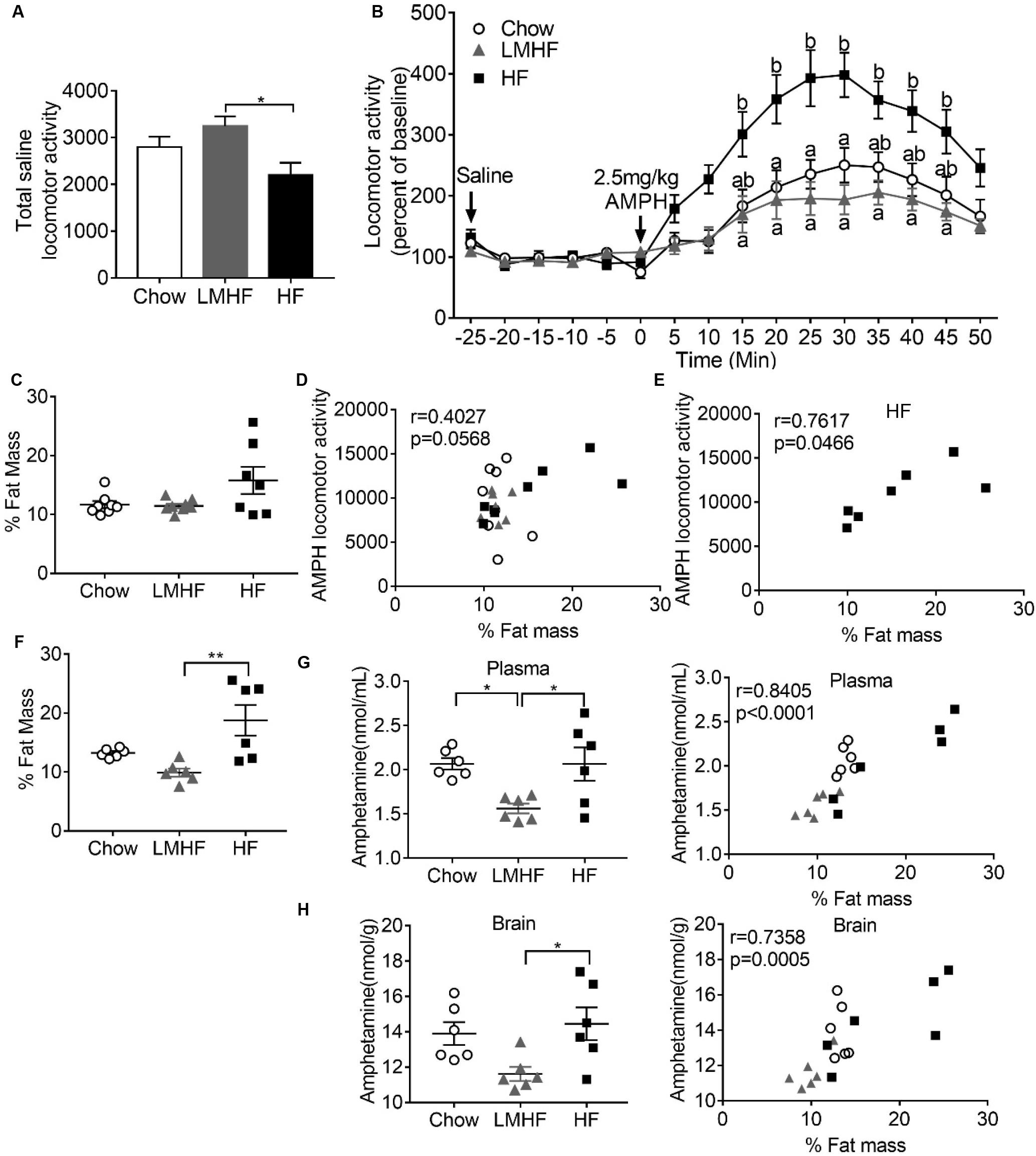
Effect of HF diet and adiposity on AMPH-induced locomotor activity and AMPH levels in plasma and brain. (A) The baseline locomotor activity was lower in WT mice fed the HF diet (HF), compared to those that were given limited access to HF diet (LMHF). n=8/group. (B) The normalized locomotor response to AMPH is higher in HF mice compared to Chow or LMHF mice. Two-way ANOVA with RM revealed a significant effect of time (p<0.0001), a significant effect of group (p=0.0015), as well as a significant interaction between time X group (p=0.0001). Different letters indicate statistically significant differences (p<0.05) at individual time points based on Bonferroni post-hoc testing. (C) Adiposity (% Fat Mass) values for each mouse used for locomotor testing. (D) Correlation between % fat mass and AMPH locomotor activity for mice in all three groups falls just outside significance threshold (p=0.0568). (E) Limiting correlation to HF mice yields a p value (0.0466) just within the threshold for significance. (F) Adiposity (% Fat Mass) values for each mouse used for determination of amphetamine levels are shown. Adiposity of HF group is significantly greater than that of LMHF group. n=6/group. (G) Panel at left shows that plasma amphetamine levels are higher in Chow or HF groups compared to LMHF group. Right panel shows strong correlation between % fat mass and plasma amphetamine levels (p<0.0001). (H) Panel at left shows that brain amphetamine levels in HF group are higher than those in LMHF group. Panel at right shows strong correlation between % fat mass and brain amphetamine levels (p=0.0005).

Examination of the %body fat across these groups revealed that the HF group shows a wide range of values (Figure 4C) allowing us to assess if there is a correlation between adiposity and amphetamine-induced locomotor activity. Correlation analysis of mice across all three groups indicates a trend that is just outside the significance range (Figure 4D). Restricting the correlation analysis to the HF group yields a correlation that is just within the significant range. However, analysis of the effect of adiposity on amphetamine levels in plasma and brain revealed a strong correlation between these variables (Figures 4F - H). Taken together, these findings indicate that elevated adiposity elicits increased brain amphetamine levels which would account for the increased locomotor response to the initial dose of amphetamine displayed by HF mice and *Tsn* KO mice.

### Selective, constitutive deletion of Tsn from DA neurons does not alter amphetamine response or adiposity

As constitutive *Tsn* KO mice display increased brain amphetamine levels, they are not suitable for assessing whether the TN/TX complex might affect amphetamine-mediated plasticity. However, in recent studies, we have found that global, conditional deletion of *Tsn* in adulthood does not elicit increased adiposity (12). In that study, conditional deletion of *Tsn* was induced by tamoxifen treatment of mice that are homozygous for a floxed allele of *Tsn* and hemizygous for the UBC-CreERT2 allele. Thus, we reasoned that we could evaluate the role of the TN/TX complex in amphetamine-induced plasticity in the absence of the adiposity confound by using conditional *Tsn* KO mice that do not display enhanced adiposity. As midbrain DA neurons express TN and TX and are the primary site of action of amphetamine, we proceeded to check whether selective deletion of *Tsn* from DA neurons would be a suitable approach for examining the hypothesized role of the TN/TX RNase in mediating amphetamine sensitization.

To assess this possibility, we generated mice that are homozygous for the floxed *Tsn* allele, *Tsn* ^fl/fl^, and hemizygous for the DAT-Cre allele [34]. Double immunostaining for TN and TH demonstrated complete loss of TN staining in midbrain DA neurons (Figure 5A). Furthermore, as found in *Tsn* KO mice [35], conditional deletion of *Tsn* from DA neurons also induces loss of TX protein. Prior to examining the effect of this genetic manipulation on responses to amphetamine, we first confirmed that selective deletion of *Tsn* from DA neurons does not affect body weight or composition compared to Tsn ^fl/fl^ mice (Figure S1). We then evaluated the impact of selective deletion of *Tsn* from DA neurons on the locomotor response to amphetamine. Since these mice are both hemizygous for the DAT-Cre allele and homozygous for the *Tsn* floxed allele, we also tested control mice that have each of these genotypic changes separately. Comparison of the effect of amphetamine on locomotor activity in these three groups indicated that conditional deletion of *Tsn* from DA neurons does not increase the locomotor response to amphetamine above that displayed by mice that are hemizygous for the DAT-Cre allele (Figure 5B). However, these results need to be interpreted cautiously, since, consistent with the recent report by Chohan et al. [36], control mice that are hemizygous for the DAT-Cre allele exhibit a reduced response to amphetamine compared to that of the other control group, mice that are homozygous for the *Tsn* floxed allele. Given this drawback of using mice with the DAT-Cre allele to evaluate changes in the effects of amphetamine, we opted to examine the effect of global, conditional deletion of *Tsn* in adulthood on amphetamine responses using tamoxifen-induced activation of the UBC-CreERT2 allele.

**Figure 5:**
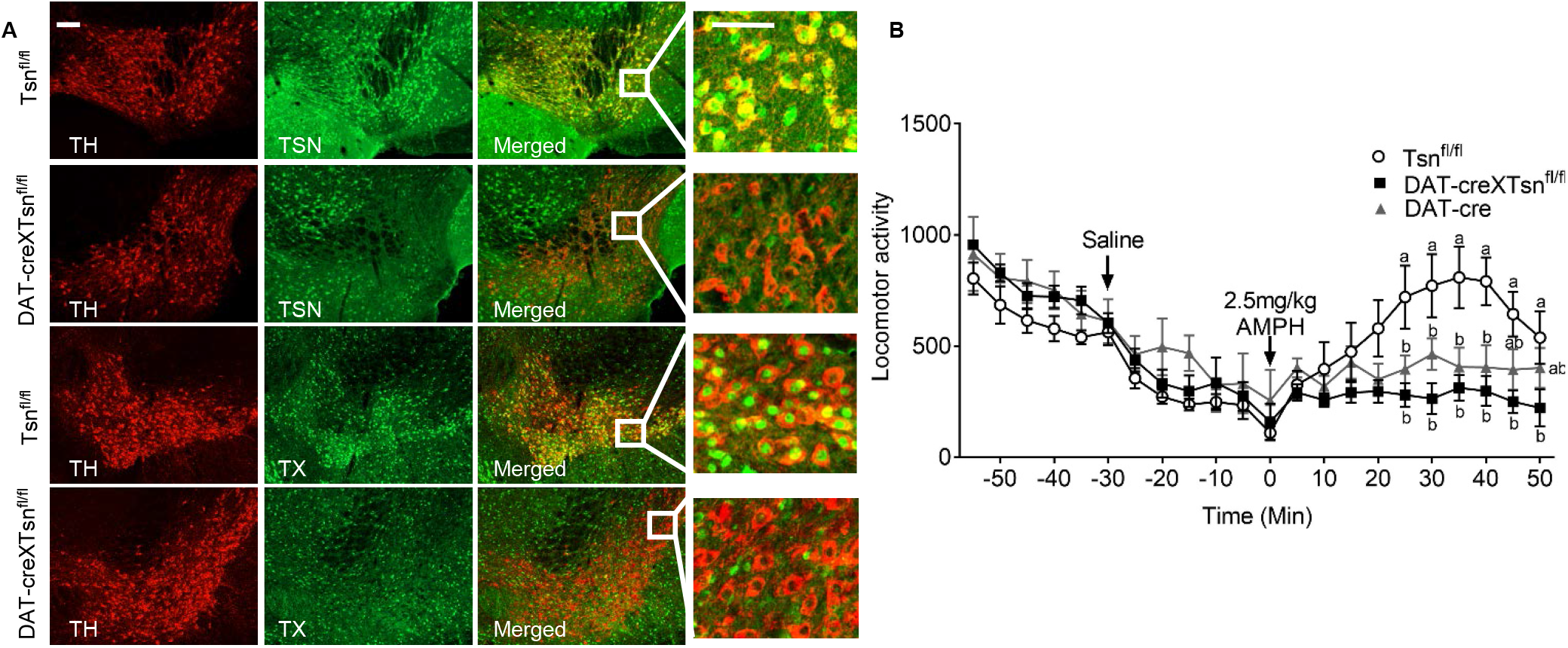
Conditional deletion of *Tsn* from DA neurons. (A) Double immunostaining of midbrain sections for TH (red) and TN or TX (green) shows that TN or TX expression was totally absent in DAT-Cre xTsn^fl/fl^ mice. Scale bar, 50 μm. (B) AMPH-induced (2.5 mg/kg i.p.) locomotor activity in DAT-Cre xTsn^fl/fl^ mice was not significantly different from the response of mice that are hemizygous for the DAT-Cre allele. n=8/group. Data are expressed as mean ± SEM. Different letters indicate statistically significant differences (p < 0.05) by two-way ANOVA with repeated measures followed by Bonferroni’s post-hoc test.

### Global, conditional deletion of Tsn in adulthood does not alter amphetamine sensitization

To check whether global, conditional deletion of *Tsn* during adulthood might affect sensitization of the locomotor response elicited by repeated administration of amphetamine, we monitored the locomotor response of *Tsn* ^fl/fl^ and UBC-CreERT2 ^+/-^/*Tsn* ^fl/fl^, that had been pre-treated with tamoxifen, to three treatments with the same dose of AMPH (2.5 mg/kg, i.p.) administered at weekly intervals. We found that *Tsn* conditional KO mice displayed decreased baseline locomotor activity prior to each administration of amphetamine (Figure 6A). Therefore, to compare the effects of amphetamine on locomotor activity between these groups, we analyzed the increase in locomotor activity induced by amphetamine relative to baseline activity which was set at 100%. Analysis of the response to each of these doses separately revealed a significant effect of time, but not of genotype (Figure 6B-D). Furthermore, the first and third doses showed a significant interaction between genotype and time. To check for an effect of genotype on sensitization, we analyzed the cumulative increase in locomotor activity elicited by amphetamine for all the doses together (Figure 6E). While this analysis revealed a significant effect of dose number, it did not find a significant effect of genotype or an interaction between dose number X genotype. Thus, we can conclude that global, conditional deletion of *Tsn* in adulthood elicits a significant increase in the locomotor response to the initial dose of amphetamine, but it does not affect amphetamine sensitization.

**Figure 6:**
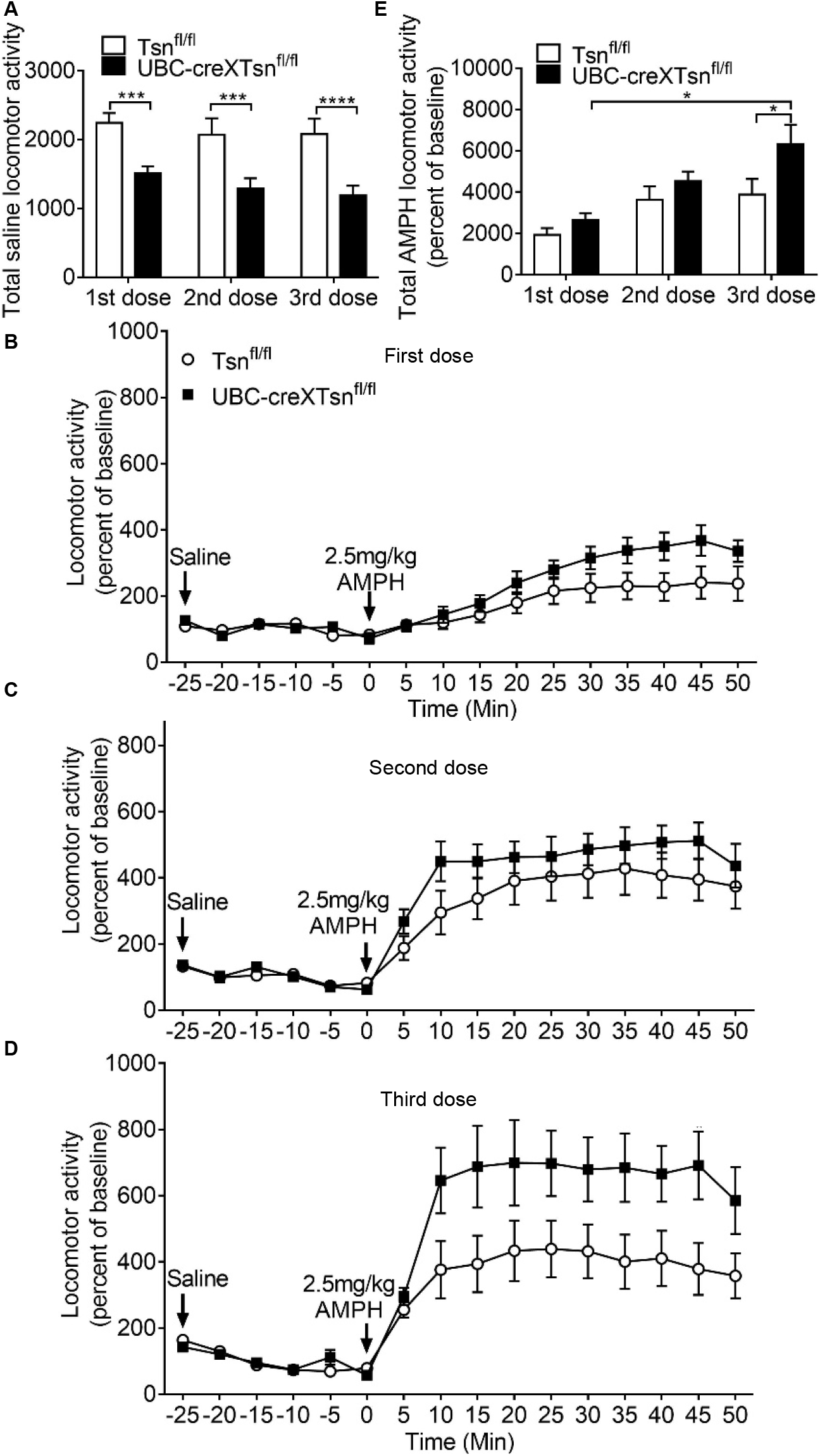
Global, conditional deletion of *Tsn* in adulthood does not affect locomotor response to AMPH. UBC-CreERT2 X Tsn ^fl/fl^ mice display lower baseline locomotor activity than Tsn ^fl/fl^ mice prior to all three treatments with AMPH (2.5 mg/kg i.p.), given at weekly intervals. UBC-Cre X Tsn^fl/fl^ mice finished tamoxifen treatment three weeks prior to amphetamine testing to induce global deletion of *Tsn*. Tsn ^fl/fl^ mice that also underwent tamoxifen treatment were used as a control group. (B-D) These panels present the time course of locomotor activity after each AMPH treatment. Two way ANOVA analysis with RM of the response to each dose separately revealed a significant effect of time with each dose, as well as a significant interaction between time X genotpe for the first (p<0.001) and third doses (p<0.001); n=8/group. Bonferroni post-hoc testing did not identify any specific time points that showed significant differences by genotype. (E) The cumulative amphetamine response elicited by all three doses were analyzed together to assess the effect of genotype on sensitization. Two way ANOVA with repeated measures revealed a significant effect of dose number (p<0.0001), but not of genotype or of dose number X genotype. Data are expressed as mean ± SEM. Statistical significance was assessed by two-way ANOVA with repeated measure followed by Bonferroni’s post-hoc analysis. *p<0.05, ***p<0.001, ****p<0.0001.

Since it is conceivable that the reduced baseline activity noted in the conditional KO mice, which were 7-8 months old, might account for the enhanced response to amphetamine, we examined their locomotor response to amphetamine at a younger age, 4-5 months old. At this age, the conditional *Tsn* KO mice and floxed *Tsn* mice displayed comparable levels of baseline activity and amphetamine still elicited increased hyperactivity in mice that had undergone conditional deletion of *Tsn* in adulthood (Figure S2).

## Discussion

While proceeding with our initial plan to examine the impact of *Tsn* deletion on amphetamine-induced sensitization, we found, unexpectedly, that the initial dose of amphetamine elicits a much stronger locomotor response in *Tsn* KO mice than WT littermates. Conceivably, this heightened response might reflect occlusion, i.e. that constitutive *Tsn* deletion might elicit a sensitized response to amphetamine even without prior exposure to amphetamine. However, prior to considering this explanation, it was important to rule out the possibility that the heightened response to amphetamine might reflect altered pharmacokinetics of amphetamine in *Tsn* KO mice. Investigating this potential mechanism seemed particularly warranted since these mice display robust adiposity, a feature that has been associated with altered pharmacokinetics of many drugs [29–31]. Furthermore, analysis of the heightened locomotor response to amphetamine of *ob/ob* mice, which display extreme obesity, revealed that these mice have higher brain levels of amphetamine [37]. By pursuing this line of research, we found that *Tsn* KO mice display higher brain levels of amphetamine providing a clear explanation for their heightened locomotor response to amphetamine.

Our working model posits that the elevated brain levels of amphetamine observed in *Tsn* KO mice are due to their robust adiposity in the context of normal body weight. As a result, these mice received a higher dose per lean body mass than WT mice, which could yield higher brain levels. Consistent with this view, we confirmed that calculating the dose of amphetamine based on lean body mass corrects the disparity in brain amphetamine levels. However, since these studies were performed with *Tsn* KO mice, they left open the possibility that the elevated brain levels of amphetamine might be secondary to other effects caused by constitutive deletion of this gene rather than increased adiposity. Therefore, to exclude this possibility we confirmed that elevated adiposity induced by switching WT mice to a HF diet also elevates brain levels of amphetamine.

Amphetamine and its analogues are widely used to treat attention deficit hyperactivity disorder (ADHD) [38, 39]. Our unexpected observation that adiposity impacts brain amphetamine levels suggests that further investigations are warranted to assess this correlation in human subjects. Furthermore, this issue is of particular concern since it is well established that ADHD is associated with obesity [40], raising the prospect that the use of standard dosing regimens in obese patients may produce higher levels of amphetamine in brain. In addition, it may be important to check if the therapeutic response or side effects produced by amphetamine may correlate with adiposity. If our findings extend to patients, then taking adiposity data into account may help determine the amphetamine dosing regimen that will optimize therapeutic efficacy.

To assess the role of *Tsn* in mediating amphetamine sensitization, we avoided the confounding effects of increased adiposity found in constitutive Tsn KO mice by switching to mice that underwent global deletion of *Tsn* in adulthood, a manipulation that does not increase adiposity. Using this approach, we were able to observe that conditional deletion of *Tsn* in adulthood does increase the locomotor response to the initial dose of amphetamine, but does not impair or enhance sensitization elicited with repeated doses. In recent studies, we have found that *Tsn* deletion impairs several acute forms of hippocampal plasticity that are dependent on rapid *de novo* protein translation triggered by neuronal stimulation, such as “spaced” LTP and synaptic tagging [8,9]. Thus, those findings indicate that these plasticity stimuli reverse microRNA-mediated translational silencing by activating the TN/TX microRNA degrading enzyme. Accordingly, it is tempting to speculate that amphetamine administration may also trigger a rapid translational response mediated by activation of the TN/TX RNase that limits the locomotor response to amphetamine, and would account for the enhanced response observed following conditional deletion of *Tsn* in adulthood. It would be interesting in future studies to test this hypothesis and examine whether TN/TX expressed in DA neurons mediates this effect.

## Acknowledgments and Financial Disclosures

This study was supported by extramural funds from NIDA (P50 DA044123; JMB), the Mid-Atlantic Nutrition Obesity Research Center (JMB), NIDA Intramural Research Program (GT and JLC), and the NIDA Medication Development Program (Z1A DA 000611; GT). None of the authors have any other financial disclosures related to the subject matter of this manuscript.

## Author Contributions

XF contributed to the design of experiments, the acquisition, analysis and interpretation of the data and writing of the manuscript. APS helped design experiments, collect and analyze data, and edit the manuscript. TMM, BL, JA, DF, MN, JLC and RR contributed to data acquisition. AS revised the manuscript. KT helped with design and interpretation of experiments. GT contributed to the design of experiments. JMB was responsible for overseeing all parts of the study and directly contributed to the interpretation of the results and writing of the manuscript. All co-authors provided final approval of the manuscript to be published.

## Supplementary Figures

**Figure S1:**
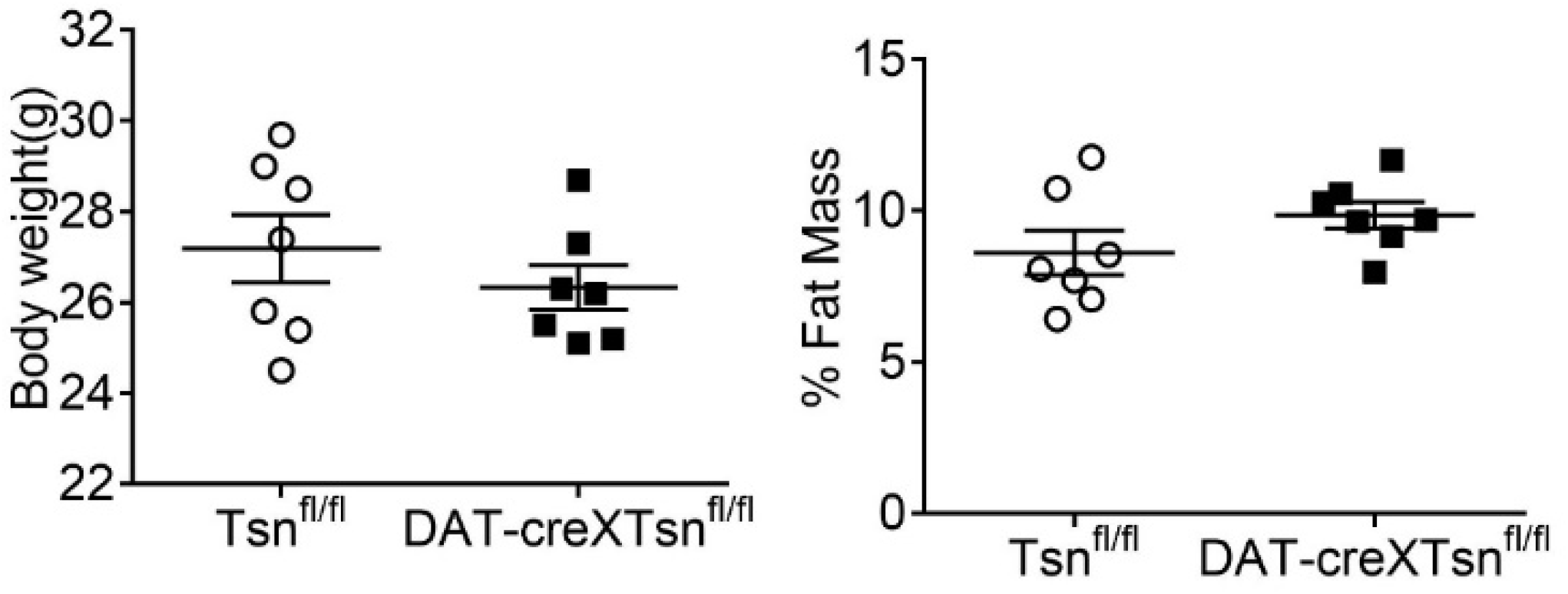
Conditional deletion of *Tsn* from DA neurons does not affect adiposity. Body weight and % fat mass were normal in DAT-Cre xTsn^fl/fl^ mice compared to Tsn ^fl/fl^ mice. n=7/group. Data are expressed as Mean ± SEM. Student’s t-test was used for statistical analysis.

**Figure S2:**
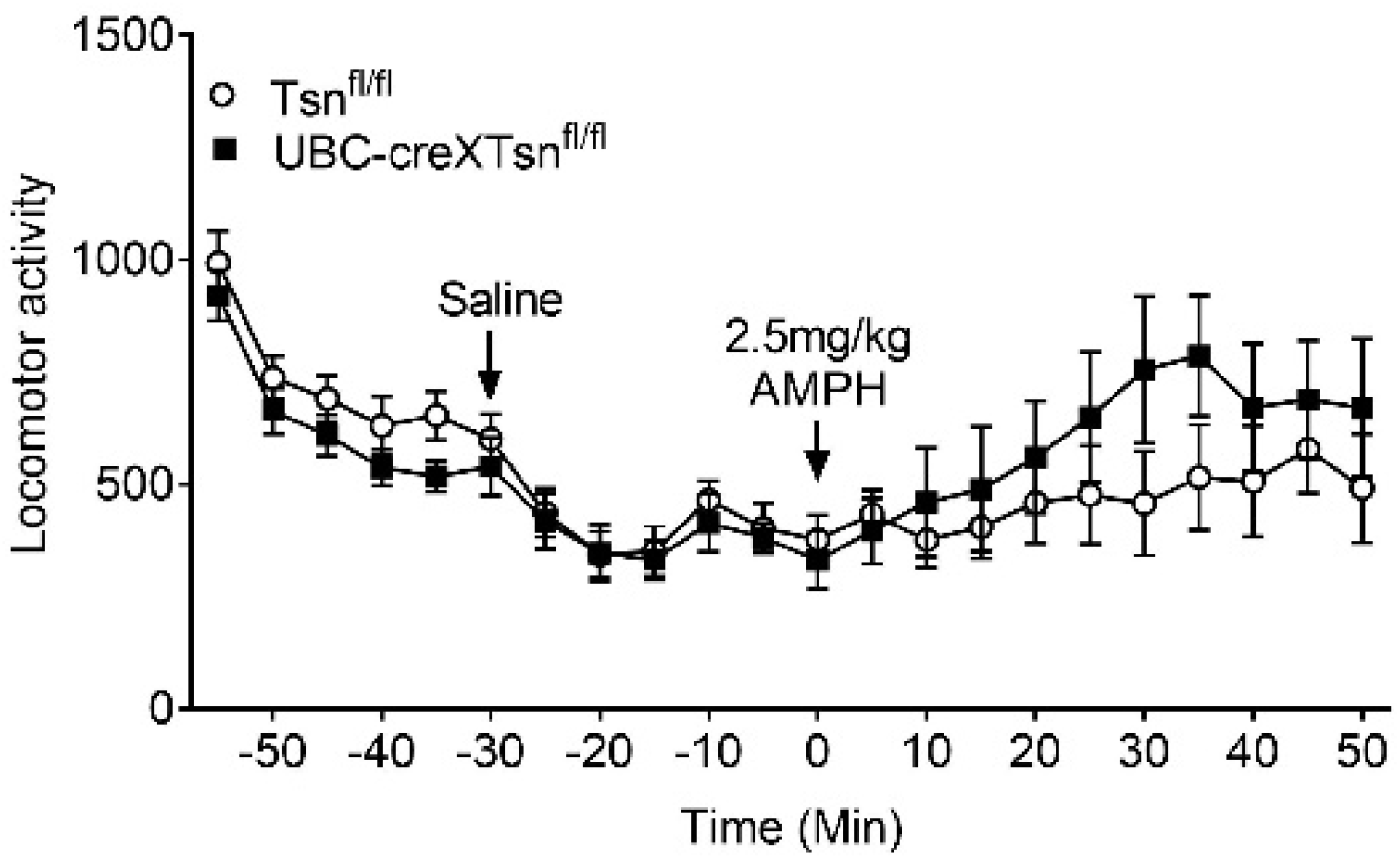
Locomotor response to AMPH in global, conditional deletion of *Tsn* mice. Two way ANOVA analysis with RM of the response to AMPH (2.5mg/kg i.p.) revealed a significant effect of time (p<0.0001), as well as a significant interaction between time X genotype (p=0.0186). Post-hoc testing did not identify any individual time points that showed significant differences by genotype. n=7/group. Data are expressed as Mean ± SEM.

